# Design: An assay based on single-polypeptide-chain heterodimeric A_2A_R/D_2_R and non-oligomerized fusions for *in vivo* analysis of their allosteric receptor-receptor interactions

**DOI:** 10.1101/065250

**Authors:** Toshio Kamiya, Takashi Masuko, Dasiel Oscar Borroto-Escuela, Haruo Okado, Hiroyasu Nakata

**Affiliations:** Department of Molecular Cell Signaling; Department of Neurology, Tokyo Metropolitan Institute for Neuroscience, *2-6 Musashidai, Fuchu, Tokyo 183-8526*, Japan; Cell Biology Laboratory, School of Pharmaceutical Sciences, *Kinki University, 3-4-1 Kowakae, Higashi-Osaka, Osaka 577-8502*, Japan; Department of Neuroscience, Karolinska Institutet, *Retzius väg8, 17177 Stockholm, Sweden;*; Neuronal Development Project, Tokyo Metropolitan Institute of Medical Science, *2-1-6 Kamikitazawa, Setagaya-ku, Tokyo 156-8506*, Japan

**Author notes:** Corresponding author;*E-mail address:* (T. Kamiya; Also, E-mail address at Showa University). Present address: Division of Gene Regulation, Institute for Advanced Medical Research, Keio University School of Medicine, Keio University, 35 Shinano-machi, Shinjuku-ku, Tokyo 160-8582, Japan (T.K.).

**Keywords:** Oligomerization, adenosine A_2A_ receptor, dopamine D_2_ receptor, receptor allostery, fusion protein, striatum, supramolecular protein assembly

## Abstract

**Background:** The adenosine A_2A_ receptor (A_2A_R) heteromerizes with the dopamine D_2_ receptor (D_2_R). In order to explore their functional interaction, we engineered previously stable single-polypeptide-chain (sc) A_2A_R/D_2L_R: whether the molecular entity of the striatal A_2A_R/D_2_R antagonism, i.e., scA_2A_R/D_2_Rs are just A_2A_R/D_2_R with the antagonism, remains unresolved.

**New Method:** To further clarify the heteromerization through the scA_2A_R/D_2L_R, we here designed supramolecularly ‘exclusive’ monomers and dimers, using the Cε2 domain of IgE-Fc or apoproteins of the bacterial light-harvesting antenna complex.

**Results:** A concept of the recptor protein assembly regulation, i.e., the selective monomer/non-obligate dimer formation was obtained. Although none of these new fusions were constructed or tested, we could aim at obtaining heterodimer-specific agents, using the scA_2A_R/D_2_R. Whether the resulting designs were explained feasibly and rationally was addressed. The structure and function of the non-obligate dimer were here discussed through scA_2A_R/D_2_R, focusing on the procedure of the membrane protein design and methods for transient protein-protein interactions.

**Summary and Outlook:** Given that upon being expressed and allosteric regulation occurs regardless of specific signal to non-specific noise (S/N) ratio, the supramolecular designs, allowing us to express selectively monomer/non-obligate dimer of class A GPCR, are experimentally testable and will be used to confirm *in vivo* that such low S/N ratio interaction between A_2A_R and D_2L_R functions in the dopamine neurotransmission in the striatum.

**Abbreviations:**

A_2A_R: adenosine A_2A_ receptor
3HA-A_2A_R: A_2A_R tagged with a triple HA epitope
Bchl: bacteriochlorophyll
BRET: bioluminescence resonance energy transfer
C: carboxy-terminal
CD: cluster of differentiation
D_2L_R and D_2S_R: the long and short form of dopamine D_2_ receptor, respectively
Fab: antigen binding fragment
Fc: Fc fragment
FcεRI: high affinity receptor for IgE
FRET: fluorescence resonance energy transfer
G_4_S: an amino acid sequence consisting of a four-glycine-repeat followed by a serine residue
GABA: γ-aminobutyric acid
GABA_B_: GABA type B receptor
GPCR: G protein-coupled receptor
G_t_: transducin
HA: hemagglutinin
HIV: human immunodeficiency virus
IC: intracellular loops
Ig: immunoglobulin
LH: light-harvesting antenna complex
mAb: monoclonal antibody
*M*_r_: molecular weight
N: amino-terminal
PD: Parkinson’s disease
PS: photosystem
RC: reaction center
Rluc: *Renilla* luciferase
sc: single-chain
TM: transmembrane
3D: three-dimensional

## 1. Introduction

In the striatal function [1,2], dopamine (DA)[3] plays a critical role. In the striatum, how the G-protein-coupled receptor (GPCR) for adenosine, subtype A_2A_ (A_2A_R)[4,5] contributes to the DA neurotransmission in the "volume transmission"/dual-transmission model [6] has been studied extensively [7]. In addition to a fusion receptor, the prototype single-polypeptide-chain (sc) heterodimeric A_2A_R/D_2_R complex (the GPCR for DA, subtype D_2_; [8])[9], several types were created and tested experimentally [10], while referring to a relationship between nanoscale surface curvature and surface-bound protein oligomerization [11-12]. In order to further explore *in vivo*, we have thus designed a new molecular tool, supermolecule scA_2A_R/D_2_R. Here, even if no experiments on their expression will be done, and thus apart from discussing based on the obtained results, the structure and function of the non-obligate dimer will deserve to be discussed through scA_2A_R/D_2_R, focusing on the procedure of the membrane protein design and methods for transient protein-protein interactions [14,15], which have been rapidly progressing and burgeoning. scA_2A_R/D_2_R is intended to draw attention to possible developments in this field and for opinion pieces in a range of alternative solutions to controversial evaluation of data obtained by various methods that so far, have been extensively reviewed elsewhere [16,17].

## 2. Perspectives

### 2.1. A trial targeting insufficient class A GPCR dimer

By a model of receptor-receptor interaction, a functional antagonistic interaction between A_2A_R and D_2_R to modulate DAergic activity has been explained [18-21]. While allostery in a GPCR hetero-dimer has been demonstrated [22-26], these class A GPCR dimers, unlike other class GPCRs, are not fully formed [27], but depend on the equilibrium between monomers and dimers [16]. Also, the recent BRET evidence indicates that class A GPCRs dissociate, merely by "being in close proximity by chance (a.k.a. "bystander BRET")"[28], and additional evidence provides a case against invariant class A GPCR oligomerization [29], but they do not mean that such insufficient class A GPCR dimers or oligomers cannot function *in vivo*. Some types of protein-protein interactions, transient or weaker, "will be found to play an even more important role" in the cells [14,15].

Interestingly, in the phase of rational design and screening, fusing the two receptors was found to stimulate the receptor dimer formation [9,10]: In the study, by fusing the cytoplasmic carboxy-terminus (C-ter) of the human brain-type A_2A_R (that is derived from Dr. Shine's cDNA [30])(**Fig. S1**) via the transmembrane domain (TM) of a type II TM protein (Section 3.3) with the extracellular amino-terminus (N-ter) of D_2_R in tandem, we created successful designs for a fusion receptor, single-polypeptide-chain (sc) heterodimeric GPCR complex A_2A_R/D_2_R, as a solution [9,10]. However, the resulting prototype scA_2A_R/D_2L_R has compact folding, i.e., fixed stoichiometry (the apparent ratios of A_2A_R to D_2_R binding sites), A_2A_R : D_2_R = 10:3 = 3~4:1 ([10]: **Graphical abstract**) and the scA_2A_R/D_2_R expression system shows that the various designed types of functional A_2A_R/D_2_R exist even in living cells, but thereby with no apparent allostery as a whole. Thus, to further clarify the heteromerization through scA_2A_R/D_2L_R, we tried to design other fusion proteins so as not to be formed/expressed as higher-order-oligomers, we called these 'exclusive' monomers (Fig. 1B/2) or dimers (Fig. 3): GPCRs are provided with general features of TM helix 3 as a structural (at tilt-angle of 35°)/functional hub and TM helix 6 moving along 14 Å after activation [31]. Taking note of A_2A_R with the bundle width diameter of about 3.6 nm (Fig. 2A), thus, using a partner to space out [32] without identifying and specifically blocking the interacting portions between the receptors, we created the designs for non-oligomerized ‘exclusive’ monomeric A_2A_R and/or D_2_R in order to exclude their dimer/oligomer formation (Fig. 1B/2)(Sections 3.1.1 and 3.1.2), and for the ‘exclusive’ dimer (Fig. 3)(Section 3.2). Such molecular self-assembled architecture will entirely hold either a monomeric receptor or single dimeric A_2A_R/D_2_R alone, but none of the oligomers.

**Fig. 1.**
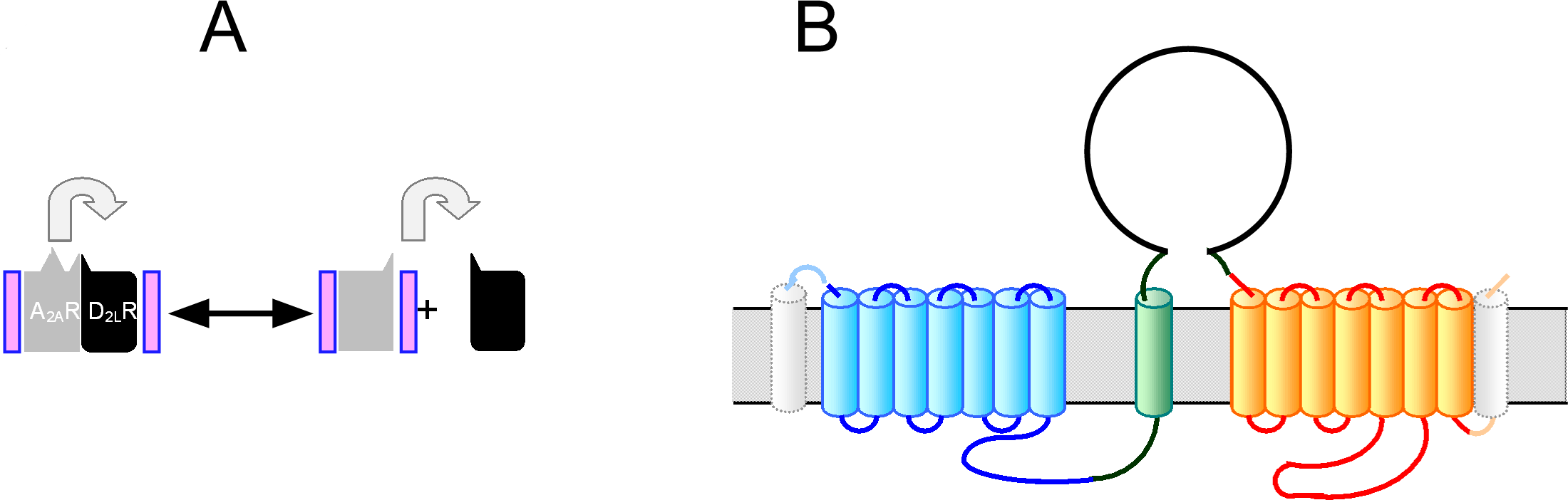
An ‘exclusive’ monomeric GPCR with the human Cε2 domain. See text in detail. The figure art (**B**) is shown which was drawn, being adapted from the published figure, i.e., a figure art (lower left) of A_2A_R-*odr4*TM-D_2L_R in Fig. 1A (pp. 140) in our previous report [9], with a written permission by the copyright owner, the Japan Society for Cell Biology.

Although none of these new fusions were constructed or tested, we could aim at obtaining heterodimer-specific agents, using the fusion receptor, scA_2A_R/D_2_R. Indeed, a universal influenza vaccine was engineered through fusing two polypeptides that originally resulted from the limited proteolysis of the native conformation of the hemagglutinin (HA), a trimeric membrane protein of influenza viruses, so as to be retained and stabilized [33]. Based on the above-mentioned BRET evidence [28], even if, assuming that A_2A_R and D_2L_R could dissociate, merely "being in close proximity by chance (a.k.a. "bystander BRET")", and that accordingly, the apparent ratio of A_2A_R to D_2_R binding sites was shown to be about 4 in scA_2A_R/D_2L_R possibly forced to form oligomers [10], the ratio in ‘exclusive’ forms of single-chain dimer should be 1 because of no or false interaction between the two receptors. On the other hand, even though the ratio in ‘exclusive’ forms of the single-chain dimer is 1, allosterism may be observed in the presence of the counter agonist. Furthermore, if the ratio in ‘exclusive’ forms of the single-chain dimer is not 1, it suggests that in the absence of the counter agonist, the specific interaction between the two happens, including protein folding during its biosynthesis [10]. Such 'exclusive' forms of single-chain dimers are just useful analytical tools allowing us to address this point.

Additionally, *in vivo* analysis of knock-in mice of the scA_2A_R/D_2_R in the postsynaptic striatopallidal γ-aminobutyric acid (GABA)-ergic medium spiny projection neurons (the indirect pathway) would unveil something new and unobservable, only in knock-in mice of A_2A_R and/or D_2_R, about the functional A_2A_R/D_2_R with the antagonism (**Table S1**): To illustrate this method, for clarity, let us consider in given days or a period for behavioral/locomotor activity tests of interest in adult mice: Although in the neuron, D_2_R and A_2A_R are coexpressed under the control of their respective promoters, as a negative control in addition to the wild-type mouse, for example, the A_2A_R-knockout mutant mice [34] that have been knocked-in with the gene containing the internal ribosome entry site sequence between the A_2A_R and the D_2_R under the D_2_R promoter, are also necessary in case of scA_2A_R/D_2_R expression under the D_2_R promoter [1,35,36], compared with mutant 1 (expressing supramolecular monomer A_2A_R) and mutant 2 (expressing supramolecular dimer scA_2A_R/D_2_R; A_2A_R=D_2L_R as shown in Table S1). The evidence [28] intimates to us that fine-tuning of dimerization of class A GPCR is just done at the respective transcription level as a genetic unit in the discipline of Jacob and Monod [37], and that upon being expressed and "being in close proximity by chance," allosteric regulation occurs by interaction between a class A GPCR and any class A GPCR or even other membrane proteins, regardless of the interaction [14,15,38] with specific signal to non-specific noise (S/N) ratio. Thus, to confirm *in vivo* that such low S/N ratio interaction between A_2A_R and D_2L_R functions in the DA neurotransmission in the striatum, we could address whether excluding such A_2A_R behavior only in postsynaptic striatopallidal GABAergic neurons, using knock-in mice with the ‘exclusive’ form of monomeric A_2A_R, results in defects compared to that of wild-type mice (**Table S1**). Also, we have already designed another type of ‘exclusive’ form of a single-chain dimer, the (core) light-harvesting antenna complex (LH) 1-based scA_2A_R/D_2L_R (Section 3.3)(T.K., unpublished).

The cytoplasmic C-ter region of A_2A_R is important for heterodimeric interaction with D_2_R [39]. Although crystallographyic analysis of GPCR hetero-dimer also remains unresolved [20], by turning from being elusive, such structural bases of these phenomena [25] are being revealed: major ligand-induced rearrangement of the TM domain interface in the metabotropic glutamate receptor homo-dimer [40], and by the single molecule analysis, multiple possible helix interfaces mediating inter-protomer associations in two functionally defined mutant luteinizing hormone receptors [41] were identified. A possible monovalent agent acting on GPCR heterodimers was referred to and a screen is needed to this end [42]. To prove the above-mentioned scGPCR-based screen as such a system is just our goal. Here, we could aim at obtaining heterodimer-specific agents [Supplemental Information (Section S3.2)], using a supramolecularly [43-45] designed fusion receptor, scA_2A_R/D_2_R, i.e., non-oligomrerized ‘exclusive’ monomer and dimer of the receptors, and *in vivo* analysis of the functional antagonistic A_2A_R/D_2_R. The possible occurrence of improper folding into a three-dimensional (3D) structure, in which the resulting receptor exhibits lower or false activity, should be avoided: This attributes to the interaction between a single ‘exclusive’ form of either receptor monomer or scA_2A_R/D_2_R and the surrounding fence-like architecture, while considering a single bond between two carbon atoms, of C-C covalent bond with a distance of 1.5 Å.

### 2.2. The molecular populations of A_2A_R/D_2L_R species in cell membranes

On the other hand, an appearance of epitopes in heteromeric A_2A_R/D_2L_R alone, but not in monomeric (and/or homodimeric) A_2A_R or D_2L_R, can occur [10], in agreement with findings on the existence of agonistic/antagonistic (active/inactive-state-specific) or dimer-specific antibodies (nanobody)(Table 1). In our case, the mouse immunization proceeded smoothly: We found many antibodies that bound either non-transfected, A_2A_R-expressed, or D_2L_R-expressed cells, as well as A_2A_R/D_2L_R-coexpressed cells, by screening hybridomas. However, the frequency of obtaining a monoclonal antibody (mAb), possibly specific for heteromeric A_2A_R/D_2L_R of interest, was very low, only one out of more than 600 wells positive upon being screened, within a total of 960 wells plated after cell fusion {[10]-(s48);(T.K., T.M., unpublished data)}. It was reported on a broad and extremely potent human immunodeficiency virus (HIV)-specific mAb, termed 35O22, which binds the gp41−gp120 interface of the viral envelope glycoprotein (trimer of gp41-gp120 heterodimers)[46](Table 1). This dimer-specific mAb was obtained despite not being immunized. The existence of virus-neutralizing mAbs, such as 35O22 that recognizes HIV-1 gp41-gp120 interface [46] and 2D22 locking the dimeric envelope proteins of dengue virus type 2 [47], is suggestive of that of heterodimer-specific mAb of our interest. Thus, the expression of homogeneous molecular species, either monomer or dimer, but not the mixture of both, followed by their membrane preparation appropriate to our needs, is necessary and worthy to be addressed experimentally. For our plan, it is advisable to start the construction of the exclusive forms of scA_2A_R/D_2L_R.

**Table 1.**
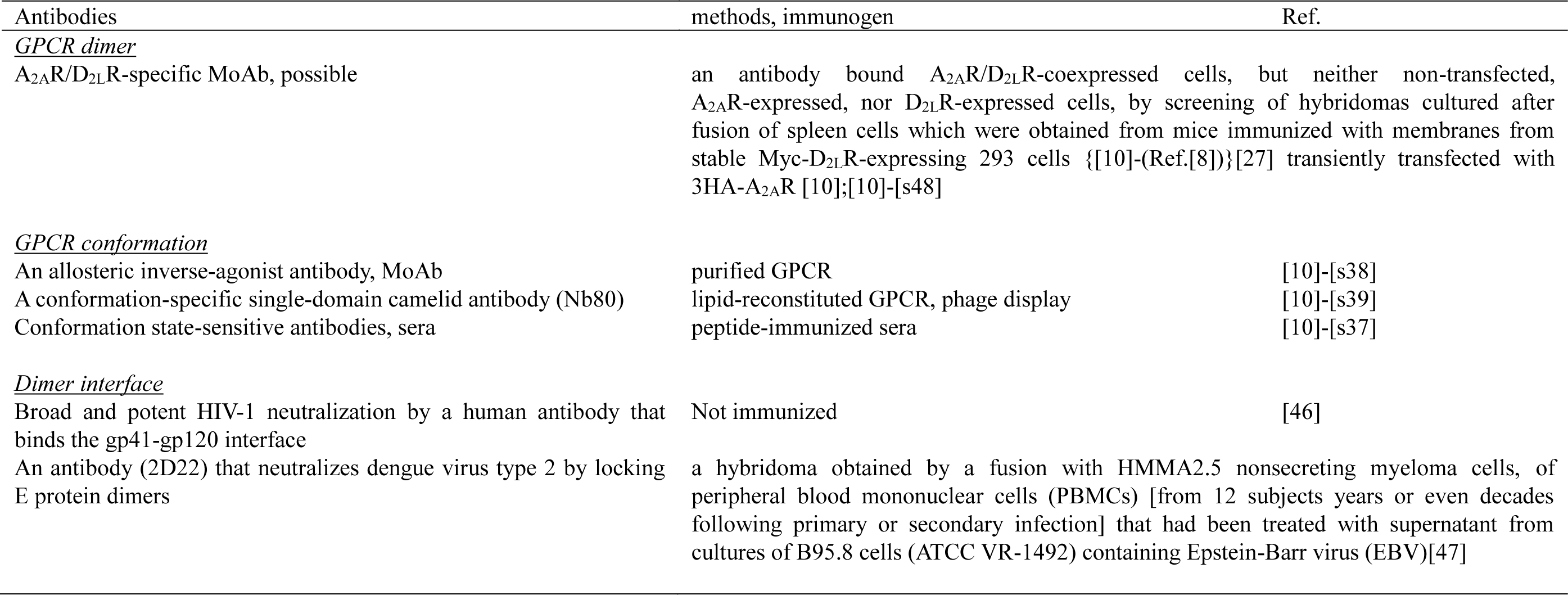
Making monoclonal antibodies specific for heteromeric A_2A_R/D_2L_R^a, Main text:^**Table Footnote**)

## 3. Architecture

### 3.1.1. Thinking about protein assembly regulation and supramolecularly designed ‘exclusive’ monomers, using the Cε2 domain of IgE-Fc

The molecular entity of allosteric modulation of A_2A_R/D_2_R remains unresolved. In order to circumvent the insufficient dimer formation of A_2A_R/D_2L_R, various scA_2A_R/D_2_R constructs, with spacers between the two receptors, were created ([10]: **Graphical abstract**): Successful design of fusions, A_2A_R-D_2_R(ΔTM1) (not shown), D_2_R-A_2A_R(ΔTM1) (not shown); the prototype A_2A_R-*odr4*TM-D_2_R; the same configuration as the prototype, but with different spacers, and the same configuration as the prototype, but with different TM (A_2A_R-TM-D_2_R); and the reverse configuration, D_2_R-*odr4*TM-A_2A_R. Using whole cell binding assays, constructs were examined for their binding activity. Although the apparent ratio of A_2A_R to D_2_R binding sites should be 1, neither was, suggesting compact folding, i.e., A_2A_R: D_2_R = 10:3 = 3~4:1; Counter agonist-independent binding cooperativity occurred in context of scA_2A_R/D_2_R. The A_2A_R/D_2L_R dimer (Fig. 1A) could show the cooperativity through the direct interaction in the presence or absence of a counter-agonist on the cell surface membrane (left). However, the monomer (here A_2A_R in gray) also could do it exclusively, not through the direct interaction between the receptors, but through the third molecules (in violet), including allosteric modulators, such as a lipid oleamide on the serotonin receptor [48-50], where a counter-agonist could induce the binding between a receptor (here A_2A_R) and the modulators, and thus make another receptor (here D_2L_R, in black) lack the same modulators, resulting in a decrease in the D_2_R affinity (i.e., the negative cooperativity, right)[23]. Another expalnation is possible (Section S3.3). This study into the bulky complexes needs to pay attention to the following matter: The cell surface on the intact living cells is, although not completely, free of curvature stress, in contrast to the intact intracellular vesicles and the membrane preparation. The membranes prepared only by homogenization are more than 10 μm in length. In preparing by sonication, they vary small unilamellar liposomes that are smaller than 50 nm in diameter, thus being highly fusogenic, to the large multilamellar ones such as only homogenates [11]. Also, note A_2A_R has a bundle width diameter of about 3.6 nm (Fig. 2). Inspired by two papers [51,52], we also designed the following fusions (Fig. 1B): First, the change between a single- and two-antigens binding by the designed antibody was reported using a hinge/domain. Then, in models of viral fusogenic proteins, both steric hindrance and conformational changes, i.e., negative cooperativity, were referred to. Accordingly, while considering the structure of the complete Fc fragment (Fc) of immunoglobulin (Ig) E, including the Cε2 domains, which is compact, bent conformation ([53]: pp. 205, the right column-line 2 from the bottom; [54]){The human IgE, as well as IgM, lacks a flexible hinge region, unlike other class/subclasses, such as IgG1, instead has a rigid Cε2 domain of Fc portion, allowing that upon the binding of an allergen to the IgE that is already bound to the high affinity receptor FcεRI, the antigen binding fragment (Fab) portion transduces it to the FcεRI [54]}, for the correct protein folding of Cε2, it was extracellularly expressed here. Thus, the expression of the ‘exclusive’ monomeric GPCRs with the human Cε2 domain (here in a loop), i.e., the C-ter, but not the N-ter, of the *odr4*TM of the prototype scA_2A_R/D_2L_R fused to the N-ter of Cε2, and its C-ter fused to the N-ter of the D_2L_R, could separate from each other. Whereas the prototype A_2A_R-*odr4*TM-D_2_R stimulates the dimerization of A_2A_R and D_2_R, this type of Cε2-intervening scA_2A_R/D_2_R makes both receptors repel each other, resulting in two 'exclusive' monomers. To this end, moreover, additional bulky molecules (gray dotted line) at both the N-ter and the C-ter of a fusion would be preferable.

**Fig. 2.**
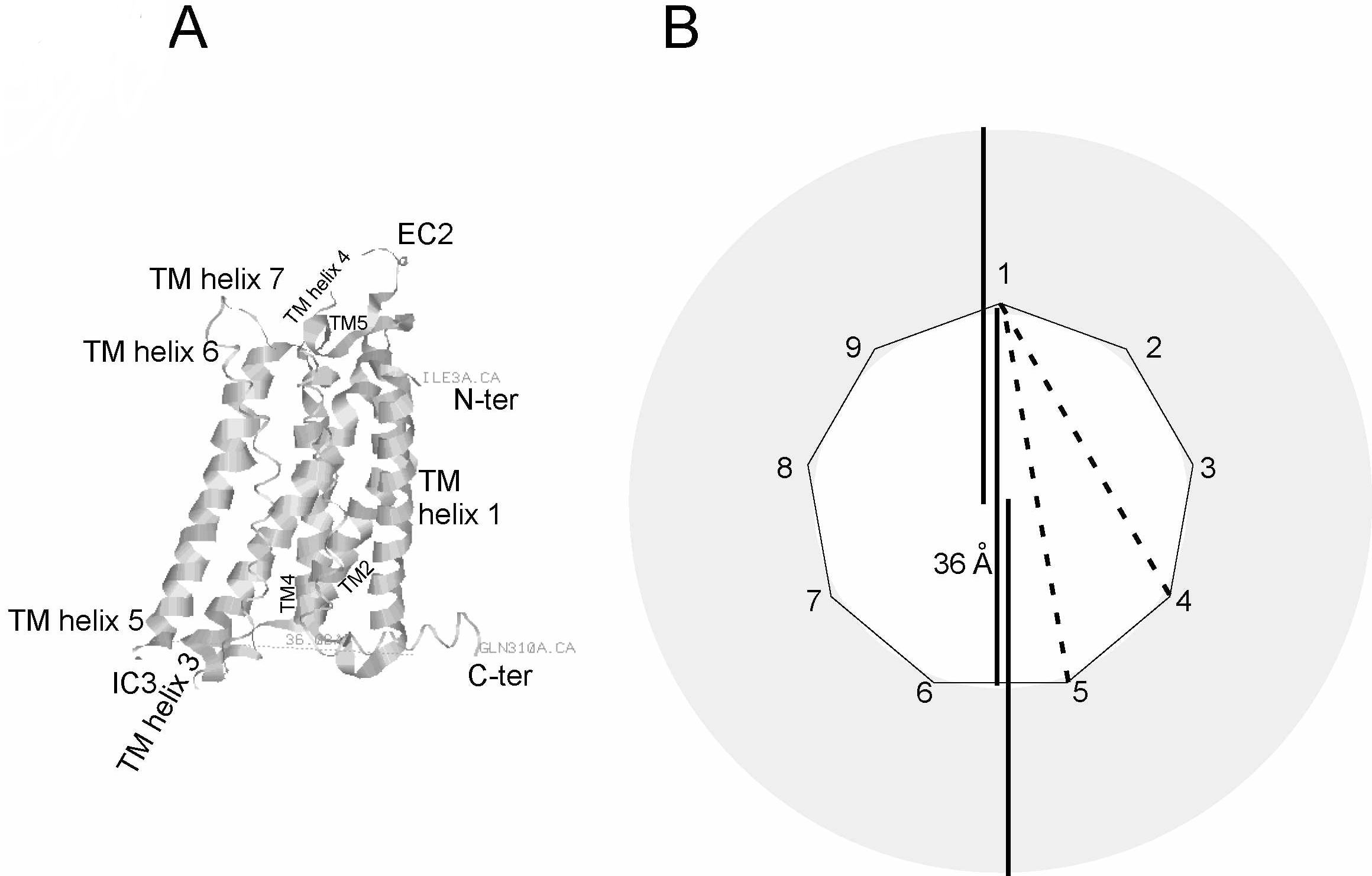
Another ‘exclusive’ monomeric GPCR with the oil-fence-like structure. (**A**) The human A_2A_R structure PDB 3EML [55] is shown with a bundle width diameter of about 36 Å (Section S2). (**B**) The light-harvesting antenna complex (LH2, gray circle) from the *Rhodopseudomonas acidophila* strain 10050 consists of 9 mer of the α-apoprotein [53 amino acids, with the intracellular N-terminus (N-ter); a nonagon with each angle numbered] packed side by side to form a hollow cylinder with a radius of 18 Å and the 9 helical β-apoprotein (41 amino acids, with the intracellular N-ter) of an outer cylinder with a radius of 34 Å, together with porphyrin-like, light-absorbing pigments bacteriochlorophyll *a*(Bchl *a*) and carotenoids [57]. Lines for 36 Å are shown in black. The expression of an ‘exclusive’ monomeric GPCR with the oil-fence-like structure, i.e., the C-ter of this α-apoprotein (Angle 1) fused to the N-ter of TM helix 1 of GPCR, and its C-ter of TM helix 7 fused to the N-ter of another α-apoprotein (between Angle 1 and either Angle 4 or 5, a GPCR is shown by a dashed line), with other 7 wild-type α-apoproteins (respectively, Angle 2, 3, and either 5~9 or 4, 6~9), could form a complex with 9 β-apoproteins and pigments.

### 3.1.2. Another ‘exclusive’ monomeric GPCR with the oil-fence-like structure: supermolecule using apoproteins of bacterial light-harvesting antenna complex

The human A_2A_R structure PDB 3EML (Fig. 2A), determined by T4-lysozyme fusion strategy [55], where most of the IC3 (Leu209^5.70^~Ala221^6.23^: The Ballesteros-Weinstein numbering scheme is shown in superscript)[56] was replaced with lysozyme from T4 bacteriophage (not shown here) and the C-ter tail (Ala317~Ser412) was deleted to improve the likelihood of crystallization, is shown with a bundle width diameter of about 36 Å (Section S2). Recent papers consider 'the most complex designed membrane proteins contain porphyrins that catalyze transmembrane electron transfer.' [43]: The peripheral LH2 complex (a circle in gray: Fig. 2B) from the purple bacterium *Rhodopseudomonas acidophila* (*Rhodoblastus acidophilus*) strain 10050 consists of 9 mer of the α-apoprotein [53 amino acids, with the intracellular N-ter; a nonagon with each angle numbered] packed side by side to form a hollow cylinder with a radius of 18 Å and the 9 helical β-apoprotein (41 amino acids, with the intracellular N-ter) of an outer cylinder with a radius of 34 Å, together with porphyrin-like, light-absorbing pigments bacteriochlorophyll *a* (Bchl *a*) and carotenoids [57]. The arrangement of this assembly differs markedly from that in the plant light-harvesting complex, where the gross assembly is formed from a <u>trimer</u> of complexes. Lines for 36 Å are shown in black. The expression of an ‘exclusive’ monomeric GPCR with the oil-fence-like structure, i.e., the C-ter of this α-apoprotein (Angle 1) fused to the N-ter of TM helix 1 of GPCR, and its C-ter of TM helix 7 fused to the N-ter of another α-apoprotein (between Angle 1 and either Angle 4 or 5, a GPCR is shown by dashed lines), with other 7 wild-type α-apoproteins (respectively, Angle 2, 3, and either 5~9 or 4, 6~9), could form a complex with 9 β-apoproteins and pigments, whereas it could not do two complexes where two α-apoproteins (Angles 1 and 4 or 5) separately interact with the respective other 8 wild-type α-apoproteins and sterically only 8 β-apoproteins, thus allowing it to be unstable probably due to the lack of a β-apoprotein. Both Bchl and carotenoids are well known to be incorporated into mammalian cells, for example, the photosensitized Pd-Bchl derivative such as WST11 as an anti-cancer drug and carotenoid vitamin A.

### 3.2. An ‘exclusive’ dimeric GPCR with the oil-fence-like structure: supramolecular dimer

Using the LH complex from *Rhodopseudomonas acidophila* strain 10050, the C-ter of an α-apoprotein (Angle 1, gray dotted line) fused to the N-ter of TM helix 1 of the human prototype scA_2A_R/D_2L_R, i.e., A_2A_R*odr4*TM-D_2L_R (colored), and its C-ter of TM helix 7 fused to the N-ter of another α-apoprotein (a given Angle of another hemi-complex, gray dotted line), are shown (Fig. 3A). Its expression of a fusion, with the remaining ~16 wild-type α-apoproteins (presumed to be 18 mer originally in total), could form a unified complex, although bending, with ~18 β-apoproteins in total and pigments, thus surrounding scA_2A_R/D_2L_R. Alternatively, the expression of an ‘exclusive’ dimeric GPCR with the oil-fence-like structure (here shown in a bending dashed line in black: Fig. 3B) could form two complexes, where two α-apoproteins (Angles 1 and 1') separately interact with the respective other ~8 wild-type α-apoproteins and ~9 β-apoproteins, thus being possibly unstable due to the lack of apoproteins. To solve this ambiguity, two units of the hetero-9-meric-αβ-apoprotein-containing LH2 complex may well be replaced by another 'fence'-like complex packed side by side to form a hollow cylinder with a larger radius, allowing them to sufficiently and stably surround a core scA_2A_R/D_2_R, as an ‘exclusive’ dimer (Section 3.3)(T.K., unpublished). Lines for 36 Å are shown in black.

**Fig. 3.**
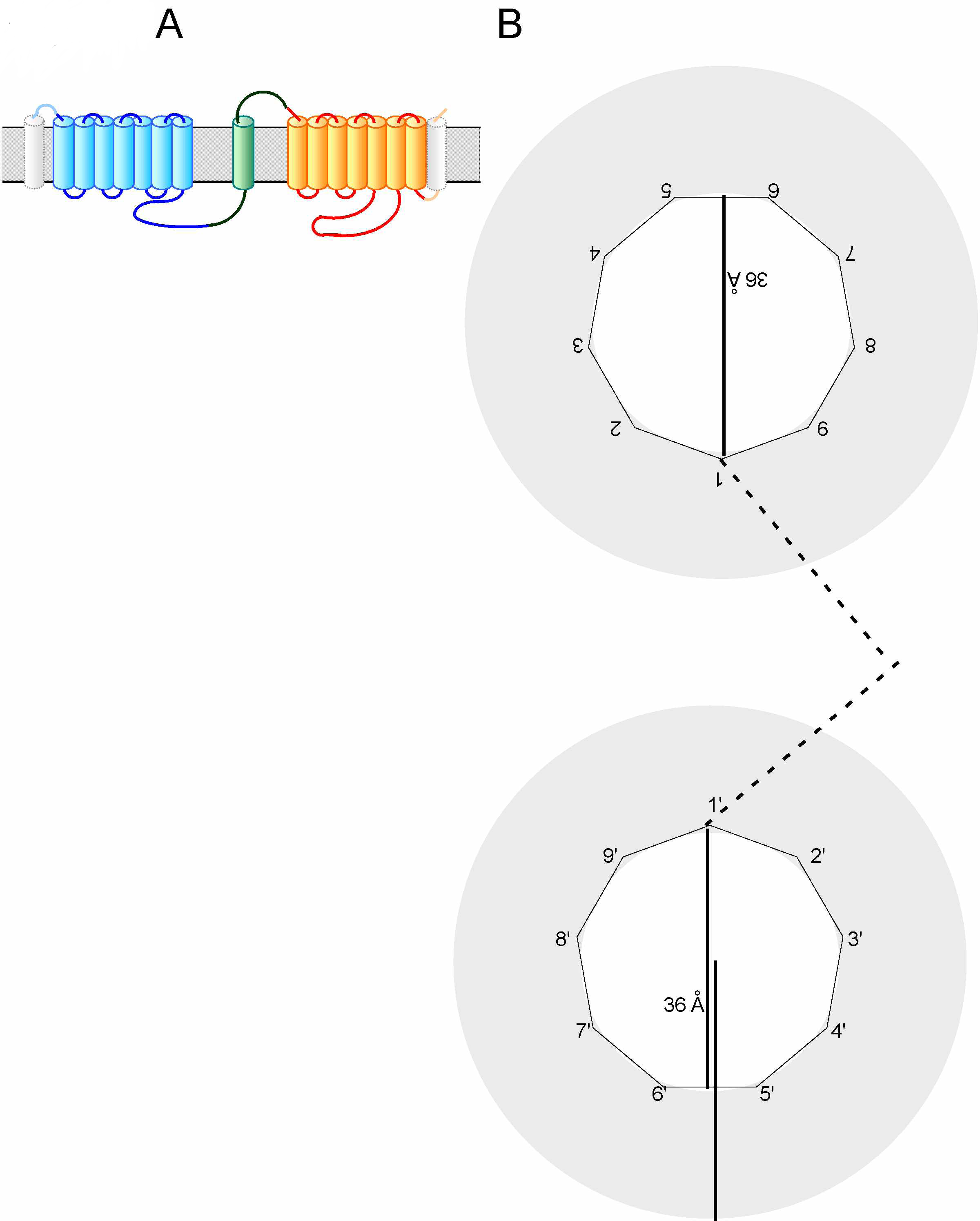
An ‘exclusive’ dimeric GPCR with the oil-fence-like structure. See text in detail. Lines for 36 Å are shown in black. The figure art (**A**) is shown which was drawn, being adapted from the published figure, i.e., a figure art (lower left) of A_2A_R-*odr4*TM-D_2L_R in Fig. 1A (pp. 140) in our previous report [9], with a written permission by the copyright owner, the Japan Society for Cell Biology. This adaptation is only a small modification as a figure art, but biologically with very important intellectual revision introduced.

### 3.3. Protein-protein interactions affect the function

Imprinted-polymers-tailor-made mimics of the 'fence'-like architecture in our supermolecule ‘exclusive’ scA_2A_R/D_2_R would be possible in the future, based on polymer chemistry [58-61]. Also, the very recent rational *de novo* protein design [62-64] would be applicable to membrane proteins. Additionally, an approach different from the 'fence'-like architecture in our supermolecule to this end would be necessary for scA_2A_R/D_2_R.

Protein-protein interactions can be classified on the basis of their binding affinities [14,15,65,66]: By definition, unlike permanent interactions with high affinities (*K*_d_ in the nM range), proteins interacting in a transient manner, either weakly or strongly, show a fast bound-unbound equilibrium with *K*_d_ values typically the μM range or less. As an example for strong transient interactions, G protein, consisting of three protomers/subunits (α, β and γ), is well known [67-70](Flock et al., 2015; DeVree et al., 2016)[71,72]. Among class A GPCRs, photoexcited rhodopsin, where i.e., "absorption of a photon causes isomerization of 11-*cis*-retinal to all-*trans*-retinal," leading to a conformational change of rhodopsin, catalyzes G protein (transducin, G_t_) cycle between the active GTP- and inactive GDP-bound states by allosterically stabilizing this transition state: Photoexcited rhodopsin has a high affinity for nucleotide-free G_t_ (Section S3.3) but not for G_t_-GDP or G_t_-GTP. Sternweis and Robishaw reported the high affinity binding of a non-hydrolyzable GTP analog, GTPγS, for G-protein in bovine brain membranes, with 0.37 nmol/mg of protein [73,74]. "Kinetic data on G protein activation are difficult to obtain,"[69], however, and parameters on the affinity between a GPCR and a G-protein have not clearly been reported so far, especially in cells [69], except for rhodopsin with *K*_d_ of nM ranges determined by cell-free, surface plasmon-waveguide resonance spectroscopy [75,76] and for a class A GPCR with *K*_d_ of around 100 nM by microscale thermophoresis [77]. A recent study, by double electron-electron resonance spectroscopy through paramagnetic labels carrying unpaired electrons that can be introduced commonly as nitroxides via usually sulfhydryl group of cysteine residues of proteins of interest [78], indicates "a minority population exhibiting substantial domain separation" between the GTPase and helical domains of G-protein α [79], although "no crystal structure of a nucleotide-bound G protein has captured a domain-separated conformation—perhaps because such conformations are less populated and less amenable to crystallization than one with the domains in tight contact—"[80]. As an example for weak transient interactions with *K*_d_ of the μM range, a complex in the plant photosynthesis, of an electron carrier protein, ferredoxin, and its dependent oxidoreductase (flavine-adenine dinucleotide flavoenzyme) has been investigated: Some bacteria and algae possess another flavoprotein, flavine-mononucleotide-containing flavodoxin. Although the ferredoxin and flavodoxin lack structural homology, interestingly, X-ray crystallography and NMR shows the molecular association of the flavoenzyme with one of these two electron carrier proteins at the same area [81-85]. Double electron-electron resonance spectroscopy and NMR chemical shift perturbation experiments [86] were also applied to analyze the intra-TM interactions of GPCR during the activation [87]. On the other hand, as for the interactions between two class A GPCRs, in addition to Ahnert *et al.*'s periodic table of protein complexes [88], the quaternary structure topologies of a tetramer of class A GPCR can be generated [20]. Recently, a paper [89] suggested that rhodopsin dimerization is possibly critical for the formation of a stable rhodopsin-G_t_ complex and the stabilization of rhodopsin pigment structure, but not for G protein-dependent responses (G_t_ activation rates), using synthetic peptides derived from bovine rhodopsin TM regions to inhibit rhodopsin dimer formation: TM4 had the greatest affinity to the rhodopsin, with a calculated apparent *K*_d_ of around 0.5 μM, and the TM1, TM2, TM4, and TM5 mixture was able to reduce the dimer from only ~65 to 40%, meaning that the interaction between two rhodopsins is stronger than this *K*_d_ value. In this context, a supermolecule scA_2A_R/D_2_R system ‘exclusive’ monomer could inhibit dimer formation completely. In addition to the superiority, another type of ‘exclusive’ dimer could also form a dimer of A_2A_R and D_2_R completely, assuming a bound-state-directed equilibrium of a core scA_2A_R/D_2_R within the oil-fence, with (1 molecule)/[π(3.6 nm/2)^2^] or 2/[π(3.6 nm)^2^] ≈ 0.1 or 0.05 nm^−2^ = 1 or 0.5 × 10^5^ μm^−2^, comparable to the concentration of rhodopsin (dimer) in the rod outer segment of 3 mM, which corresponds to a surface density of 27,000 μm^−2^ [67].

In our previous report [9,10], among GPCR protein-protein interactions, such as the disulfide bond formation of the N-ter, coiled-coil interaction of the cytoplasmic C-ter, and TM interaction [90], <u>a type II TM protein with a cytoplasmic N-ter segment, single TM, and extracellular C-ter tail</u>, i.e., the *Caenorhabditis elegans* accessory protein of odorant receptor (*odr4*)[91] was first selected for connection between the N-ter receptor half (A_2A_R) and the C-ter receptor half (D_2L_R, the long form of D_2_R) of the scA_2A_R/D_2L_R. Then, it was demonstrated that an insertion of some other TM sequence instead of the *odr4*TM sequence (TILITIALIIGLLASIIYFTVA) works similarly, or that it does not have to be *odr4*TM to work, using the scA_2A_R/D_2L_R designed with another TM of a type II TM protein, the human low affinity receptor for IgE designated CD23 (3.1).

Human ortholog, c1orf27 (chromosome 1 open reading frame 27: NP_060317.3; 454 aa)(http://www.ncbi.nlm.nih.gov), which was originally identified as human ODR4 and shown to have sequence, 432-IgViaAftVAVLAagIsFhyf-452 (Regions representing identity at that position or conserved amino acids to ODR4 TM are indicated in capital letters: [92]), has been regarded as the ODR4-like (pfam14778) conserved protein domain family. Smart blast search of *odr4*TM gave 3 hits, respectively, with E-values of 0.002, 0.003, and 0.012: hypothetical protein NOC27_163 (EDZ66836.1, *Nitrosococcus oceani* AFC27), photosystem (PS) II reaction center (RC) protein ycf12 (YP_009056868.1, *Bathycoccus prasinos*), and hypothetical protein MXAN_3681 (ABF89922.1, *Myxococcus xanthus* DK 1622).

Among these hits, a hit of a photosynthetic RC protein is impressive to us because of a use of its surrounding apoproteins in the bacterial LH complex in the supramolecularly designed scA_2A_R/D_2_R as described above. Blastp search of *odr4*TM against all non-redundant GenBank databases and the human genome gave no hits for human CD23.

On the other hand, CD23 is highly conserved among mammals. Smart blast search of human CD23TM (TQIVLLGLVTAALWAGLLTLLLLWHW) gave 2 additional hits, respectively, with E-values of 0.003 and 0.028: sulfonamide resistance protein (CRH97060.1, *Streptococcus pneumonia*) and 4-hydroxybenzoate octaprenyltransferase (WP_037755226.1, *Streptomyces sp*. AA0539).

Blastp search of human CD23TM against genomes of *Drosophila melanogaster*, *Saccharomyces cerevisiae*, and *C. elegans* gave no hits with E-values fewer than 0.01. This suggests that human CD23TM had been selected as a preferable experimental design due to little relationship between the two protein TMs, while only the structures of the extracellular region of the human CD23 were determined (PDB ID: 4KI1 [93]; 4J6J [94]; 4G96 [95]; 4EZM [96]; 2H2R [97]; 1T8C [98]).

In photosynthesis, particular protein complexes, including the PS or the LH1 and the RC [more strictly, PSI and PSII form supercomplexes with the RC and the light-harvesting proteins, i.e., respectively, PSI-type RC and LHCI, and PSII-type RC, core light-harvesting proteins (CP43 and CP47) and peripheral LHCII, in aerobic photosynthesis (algae, cyanobacteria, and plants); PS of a type RC, core LH1 and peripheral LH2 in anaerobic photosynthesis (purple bacteria)], contribute to the highly efficient, light-induced charge separation across the membrane, followed by electron transport. Whether this property is negligible or not in our 'exclusive' supramolecular forms has not been tested: rearrangement of each molecule in this architecture or exchange with similar but unrelated molecules [88] would be necessary to inhibit this photosynthetic light reaction completely and surround the prototype scA_2A_R/D_2_R.

An important finding on PSII biogenesis has been recently reported: PSII RC protein D1 is synthesized as a precursor in the *Arabidopsis* chloroplast thylakoid lumen, and cleaved by the D1 C-terminal processing enzyme (AtCtpA). Unlike its cyanobacterial counterpart, *Arabidopsis* CtpA is essential for assembling functional PSII core complexes, dimers, and PSII supercomplexes [99]. This paper shows a discrepancy in PSII protein assembly [88] between cyanobacterum and a land plant *Arabidopsis*. Thus, whether in animal cells the LH is attached to form our supermolecules, and is expressed or not, requires testing. The D1 and D2 subunits of the PS II and the M and L subunits of the bacterial photosynthetic RC are members of Photo RC (cl08220) protein superfamily. Unlike the cyanobacterium *Thermosynechococcus elongatus*, {from which the membrane-intrinsic part of homodimeric PSII comprises (in each monomer) the antenna proteins CP47 (with 6 TM helices) and CP43 (with 6 TM helices), the RC subunits D1 (with 5 TM helices) and D2 (with 5 TM helices), and 13 small subunits, among them cytochrome (cyt) b-559 with 2 TM helices [100,101],} the RC from the purple bacteria such as *Rhodopseudomonas* (*Blastochloris*) *viridis* and the thermophilic, *Thermochromatium tepidum*, consists of the four protein subunits L (with 5 TM helices), M (with 5 TM helices), H (with a TM helix) and c-type cyt (without TM)[102,103]. "The *T. tepidum* LH1−RC core complex is composed of cytochrome (Cyt), H, L and M subunits for the RC, α16β16 subunits for the LH1 and 80 cofactors" [103]. In addition, 1) based on that "When the RC was removed the ‘empty’ LH1 complex became more circular." [104], 2) and that it remains unknown what structural features lead to the consequent differences in the nonameric and octameric apoprotein assemblies in LH2, respectively, from *Rhodopseudomonas acidophila* and another purple bacterium, *Rhodospirillum* (*Phaeospirillum*) *molischianum* [105], thus meaning that it remains just unknown why LH1 surrounds the RC but LH2 does not, 3) and furthermore, due to the difficulty in membrane-protein topology prediction itself (the next paragraph), in any case up to 25% precision, "to predict interaction sites from sequence information only" {[106]: Supporting Information-[Table 9. List of PDB IDs of all monomers (Test-set 1)](pp. 38, line 27), human A_2A_R (PDB: 3EML)(Fig. 2A)(Section 3.1.2) defined as a monomer}, multiple sequence alignment was done between these proteins. In order to obtain a possible architecture that surrounds any class A GPCRs unaffectedly and that forms functional supermolecules of interest, structural similarity between class A GPCRs (A_2A_R/D_2_R) and RC proteins was not searched. Instead, for the interaction between *odr4*TM in prototype scA_2A_R/D_2_R and TMs of its surrounding (fence-like) LH proteins, multiple alignment was done between *odr4*TM and TMs of core RC proteins. The results indicate no relationship between the *C. elegans odr4*TM or human CD23 TM and TM helices of the RC subunits (L/M/H) of *T. tepidum* or *R. viridis* (**Fig. S2A/B**). It suggests that αβ subunits for the LH surrounding scA_2A_R/D_2_R do not affect their core GPCR itself, even if each protein in this supermolecule is assembled to form it. Alternative interactions between core GPCRs and surrounding LH subunits in our supermolecules remain unknown because our designs are just back-of-envelope sketches and 3D-models, especially of membrane proteins, have also their limitations [107-109].

Proteome-wide studies of membrane-protein topology have been done in an evolutionary context [110,111]: "Current topology-prediction methods are all based on the classic view of membrane-protein topology""but much work remains to be done before these more advanced types of prediction reach the level of accuracy of the simple topology-prediction schemes." Unlike cytosolic proteins, proteins transported across or integrated into the cellular membranes have the feature [110,111]. Soluble proteins synthesized as immature ones usually have the N-ter, cleavable signal sequence, which is a segment of 7~12 hydrophobic amino acids. Predicting the site of cleavage between a signal sequence and the mature exported protein is possible [112,113]. Membrane proteins have different topologies in the lipid bilayer, with one or more TM segments composed of about 20 hydrophobic amino acids; After the N-ter signal sequence is cleaved, the protein has a new N-ter extracellularly, and thus with an additional hydrophobic segment as a TM, is just a type I membrane protein.

While the N-ter hydrophilic region, without the N-ter hydrophobic signal sequence, of a protein (type II membrane protein with the intracellular N-ter) remains in the cytosol, additional hydrophobic segments function as TMs. Additionally, GPCR superfamily members have seven TM helices and a Nout−Cin orientation (that is with an extracellular N-ter and a intracellular C ter). With these points, our plan on supermolecules is complex due to the two additional LH2 α-apoprotein-fusions to, respectively, the N-ter and C-ter of the prototype A_2A_R/D_2_R, but is very interesting, experimentally testable and will help to broaden our knowledge.

## 4. Summary and Outlook

The supramolecular designs of scA_2A_R/D_2L_R, 'exclusive' monomers and dimers, using the Cε2 domain of IgE-Fc or apoproteins of the bacterial light-harvesting antenna complex, allowing us to express class A GPCR by recptor protein assembly regulation, i.e., the selective monomer/non-obligate dimer formation, are experimentally testable and will be used to confirm *in vivo* that such low S/N ratio interaction between A_2A_R and D_2L_R functions in the dopamine neurotransmission in the striatum.

## Author contributions

T.K. contributed to the planning and interpretation of all of the experiments, and conducted all of them, performed design [at The University of Tokyo General Library (Hongo, Bunkyo-ku, Tokyo)][from April 1, 2013 to date (this submission)] of vectors for the non-oligomerized ‘exclusive’ monomer and dimer, experiments such as antibody screening at TMIN (Kamiya et al., 2015), *in silico* search and wrote the paper; O.S. contributed to the interpretation of all of the experiments done at TMIN (Kamiya et al., 2015); T.M. performed immunization and cell fusion to make hybridomas at Kinki University, and with T.K., wrote the paper (Section 2.2 and **Table Footnote**). H.O. performed design (It started from December 2014) of vectors for *in vivo* analysis and with T.K., wrote the paper (in part, Section 2.1; Supplemental Information-Section S3.2). K.F. supervised the project of the electrophysiological studies (It started from November 2007)(Kamiya et al., 2015). With T.K., D.O.B.-E. analyzed the A_2A_R SNP and wrote the paper [Supplemental Information-Sections S3.1, S3.3, and S6 (Figure S1)]. H.N. supervised the overall project by the end of March 2007. H.N. also contributed to the interpretation of all of the experiments by the end of March 2007. All authors commented on the manuscript, except for H.N. who is at present under a departure from research activity.

## Acknowledgments

We thank Drs. O. Saitoh (Nagahama Institute of Bio-Science and Technology, Nagahama) and K. Yoshioka (Kanazawa University, Kanazawa) for their respective contributions (Kamiya et al., 2015), Mr. M. Woolfenden for English proofreading of a part of this manuscript, and Dr. K. Fuxe (Karolinska Institutet, Stockholm) and Dr. H. Saya (Keio University, Tokyo) for encouragement. This work was supported in part by grants for Scientific Research from the Ministry of Education, Culture, Sports, Science and Technology (MEXT) (#14657595, 13670109: to T.K., O.S., and H.N.). This work was also supported in part by grants of intramural budget (TMIN to T.K., O.S., and H.N.). T.M. was also supported by the “Academic Frontier” Project for Private Universities: matching fund subsidy from MEXT, 2005~2007. H.N. was also supported by a CREST program of the Japan Science and Technology Agency.

The authors declare no competing financial interests.

## Table Footnote

### Making monoclonal antibodies specific for heteromeric A_2A_R/D_2L_R—

This item is to discuss “the frequency of the monoclonal antibody (mAb) obtained, possibly specific for heteromeric A_2A_R/D_2L_R of interest” (Section 2.2), while in our previous report, suggesting the existence of a mAb “specific for heteromeric A_2A_R/D_2L_R (Fig. 2B); [s48];(T.K., T.M., unpublished data): we failed to establish it as a clone.” ([10])-Supplemental Information: pp. 25, line 8). In order to obtain a mAb specific for heteromeric A_2A_R/D_2L_R, partially purified A_2A_R/D_2L_R was not used as an immunogen because of being unexpectedly unable to be stored at −80 °C in prior experiments. Also, prototype scA_2A_R/D_2L_R was not used as an immunogen at that time because whether scA_2A_R/D_2L_R can simulate wild-type A_2A_R/D_2L_R remains unresolved as described [10]. Thus, mice were immunized with membranes of the coexpressed receptors. The handling of mice and all procedures performed on them were approved by the Animal Care and Use Committee of the School of Pharmaceutical Sciences, Kinki University. To check for the desired antibodies, samples of serum taken from the mice that had been immunized with a mixture of membranes from stable Myc-D_2L_R-expressing 293 cells [27] transiently transfected with 3HA-A_2A_R as the immunogen [10] and adjuvants [114], either CpG DNAs, Ribi, or TiterMax, were diluted ten- to 100-fold and assayed by an immunofluorescence-based screening method. In this method, as a set, all non-transfected, (transient) A_2A_R-expressed, (transient) D_2L_R-expressed human embryonic kidney (HEK) 293T, and (transient) 3HA-A_2A_R/(stable) Myc-D_2L_R-expressing 293 cells were plated separately directly onto 30-well glass slides [115,116] or glass coverslips, both of which had been treated with polyethylenimine-coating as usual, and fixed with 4% paraformaldehyde in Dulbecco‘s phosphate-buffered saline (D-PBS), subjected to sample incubation (anti-tag antibody was used as a control) followed by washing, and mounted, respectively, with glass coverslips or on slides using Fluoroguard^R^ [10]. The results indicated a high serum titer compared with the normal mouse serum before the immunizations begin, and that this hybridoma screening assay works well. Fusion of spleen cells, which were obtained from the immunized mice, was done as described before [117], and the cells were plated onto selection medium-filled wells of ten 96-well plates and screened. Viable hybridomas in the 958 wells survived at first (on November 8, 2003). Transient expression studies with HEK293T cells were performed as previously described [10]. On immunoblot analysis, cell membranes were subjected to SDS-PAGE and transferred to a nitrocellulose membrane as previously described [10]. Immunofluorescence studies were performed as previously described [10].

